# Robustness of time domain near infrared spectroscopy to variations in skin pigmentation

**DOI:** 10.1101/2024.07.25.605070

**Authors:** Michele Lacerenza, Caterina Amendola, Ilaria Bargigia, Alessandro Bossi, Mauro Buttafava, Valeria Calcaterra, Davide Contini, Vamshi Damagatla, Fabio Negretti, Virginia Rossi, Lorenzo Spinelli, Sara Zanelli, Gianvincenzo Zuccotti, Alessandro Torricelli

## Abstract

Recently, skin pigmentation has been shown to affect the performance of pulse oximeters and other light-based techniques like photoacoustic imaging, tissue oximetry, and continuous wave near infrared spectroscopy. Evaluating the robustness to changes in skin pigmentation is therefore essential for the proper use of optical technologies in the clinical scenario. We conducted systematic time domain near infrared spectroscopy measurements on calibrated tissue phantoms and *in vivo* on volunteers during static and dynamic (i.e., arterial occlusion) measurements. To simulate varying melanosome volume fractions in the skin, we inserted, between the target sample and the measurement probe, thin tissue phantoms made of silicone and nigrosine (skin phantoms). Additionally, we conducted an extensive measurement campaign on a large cohort of pediatric subjects, covering the full spectrum of skin pigmentation. Our findings consistently demonstrate that skin pigmentation has a negligible effect on time domain near infrared spectroscopy results, underscoring the reliability and potential of this emerging technology in diverse clinical settings.

## Introduction

Nowadays, light-based diagnostic and therapeutic devices are extensively used in the clinics^1^. Ultraviolet and blue-green (300-500 nm) light has limited penetration in biological tissue due to the combination of high absorption and scattering, therefore investigation is limited to the external part of tissues (e.g., skin or surface of exposed organs)^2^. Conversely, in the red and near infrared (NIR) part of the electromagnetic spectrum (600-1100 nm), light penetrates deeper into biological tissues and can be re-emitted at tissue boundaries thanks to the limited absorption by major tissue constituents (i.e., water, blood, lipids) and to the naturally decreasing scattering due to tissue structures^3^. Valuable information on physio-pathology can thus be derived in a noninvasive way. In the NIR spectral region, light scattering overwhelms absorption and photon propagation is typically described in the framework of the radiative transport theory with the diffusion approximation^4^. The term Diffuse Optics is currently used to refer to spectroscopy and imaging with diffusing photons. Examples of applications on humans include brain monitoring and mapping^5,6^, muscle oximetry^7,8^, optical mammography^9,10^, and perfusion monitoring^11^.

Since NIR light must travel twice through the skin to reach the target tissue (e.g., muscle, brain) and before being eventually detected at tissue boundaries, skin pigmentation, determined by the presence of melanin in the epidermis, could potentially play a role in determining the overall signal quality and characteristics. The effect of skin pigmentation (as gauged by the Fitzpatrick scale^12^ or the Monk skin tone scale^13^) on the performance of light-based devices was considered in some studies on pulse oximeters providing estimates of peripheral oxygen saturation (S_p_O_2_)^14,15^, and on continuous wave (CW) near infrared spectroscopy (NIRS) devices measuring tissue oxygen saturation (S_t_O_2_) in brain cortex or muscle^16-19^. Although not systematic, these studies suggested that skin pigmentation affects oxygen saturation measurements, particularly in subjects with dark pigmented skin^17^. However, this issue has been somehow overlooked until the COVID-19 pandemic, when several clinicians dramatically observed less accurate S_p_O_2_ readings by pulse oximeters in patients with dark skin, especially in the low saturation regime^20-23^. Thus, understanding the effect of skin pigmentation on the overall performance of pulse oximeters has rapidly become an important issue in the research community and a topic of concern in the standardization committees^24^. It is worth noting that also NIRS devices (due to the similarity with pulse oximeters^25^) and - more generally – all light-based devices might be influenced by skin pigmentation. Recent studies on the photoacoustic technique demonstrated measurement bias, including an overestimation of S_t_O_2_, in darker skin types^26,27^. Other studies on NIRS devices operating in the CW modality (i.e., with light sources emitting constant intensity and photodetectors measuring light attenuation) reported variable performance with different skin melanin content^28-30^.

In the present study, we deal with the effect of skin pigmentation on time domain (TD) NIRS devices. The TD approach^31-33^, based on the use of short laser pulses, fast photodetectors, and timing and counting electronics with picosecond time resolution, measures the distribution of time-of-flight (DTOF) of photons re-emitted from the tissue. The TD modality enables the estimate of the absolute values for the optical parameters (absorption coefficient, μ_a_; reduced scattering coefficient, μ_s_’) and of the derived hemodynamic quantities (oxyhemoglobin concentration, O_2_Hb; deoxyhemoglobin concentration, HHb; total hemoglobin concentration, tHB=O_2_Hb+HHb; tissue oxygen saturation S_t_O_2_=O_2_Hb/tHb). Furthermore, working in the TD, depth sensitivity is enhanced thanks to the fact that photons detected at later time-of-flights have a greater probability of sampling deeper regions^34^. These two characteristics result in a natural lower sensitivity of TD NIRS to changes in absorption in the very first millimeters of tissue, making it potentially insensitive to changes in skin tone^31-34^.

We conducted a comprehensive investigation using systematic TD NIRS measurements on calibrated tissue phantoms and *in vivo* on volunteers. To simulate varying melanosome volume fractions (M_f_) in the skin, we prepared thin tissue phantoms made of silicone and nigrosine (skin phantoms). These skin phantoms effectively mimicked different skin pigmentation levels. By inserting the skin phantoms between the target sample and the measurement probe, they allowed us to evaluate the robustness of TD NIRS under controlled conditions, both static and dynamic (i.e., arterial occlusion). Additionally, we performed an extensive measurement campaign on a large cohort of pediatric subjects, covering the full spectrum of skin pigmentation.

## Results

### TD NIRS static measurements with skin phantoms on bulk phantoms

Figure 1 shows the absorption and the reduced scattering spectra in the 620−1100 nm wavelength range measured with a broadband TD NIRS device on bulk solid phantoms B6 (nominal values at 800 nm: μ_a_ = 0.25 cm^-1^, μ_s_^’^ = 10 cm^-1^) and C4 (nominal values at 800 nm: μ_a_ = 0.15 cm^-1^, μ_s_^’^ = 15 cm^-1^) with different skin phantoms mimicking M_f_ = 0%, 2%, 14%, and 43% placed on top.

**Fig. 1.**
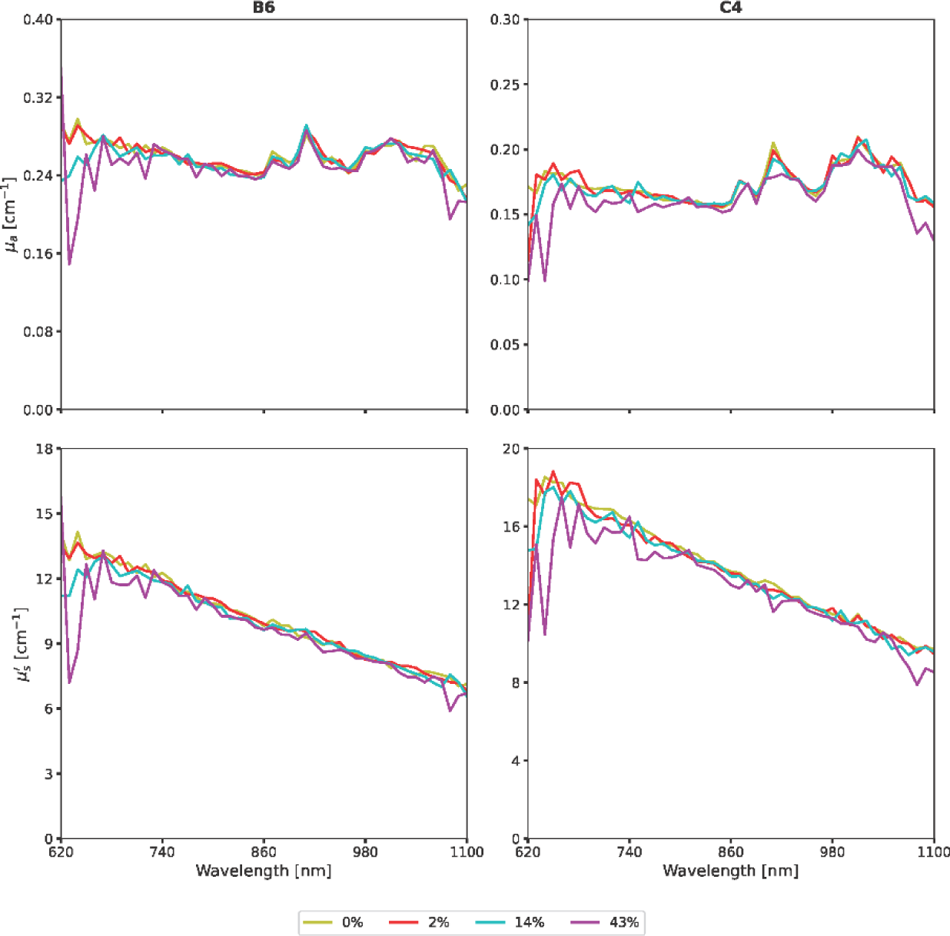
Absorption coefficient (top row) and reduced scattering coefficient (bottom row) spectra of phantom B6 (left column, nominal values at 800 nm: μ_a_ = 0.25 cm^-1^, μ ^’^ = 10 cm^-1^) and C4 (right column, nominal values at 800 nm: μ_a_ = 0.15 cm^-1^, μ ^’^ = 15 cm^-1^) with skin phantoms mimicking different skin pigmentation tones (M_f_ = 0%, 2%, 14%, 43%) placed on top.

For M_f_ = 2% and 14% μ_a_ and μ_s_’ are almost indistinguishable from the case M_f_ = 0% in both phantom B6 and phantom C4 practically at all wavelengths, with small differences only below 700 nm. For M_f_ = 43% differences can be noted below 800 nm and above 1000 nm. To better appreciate the difference between the control case M_f_ = 0% and the pigmented cases (M_f_ = 2%, 14% and 43%) as a function of the corresponding mean value, Bland-Altman plots are reported in Fig. 2.

**Fig. 2.**
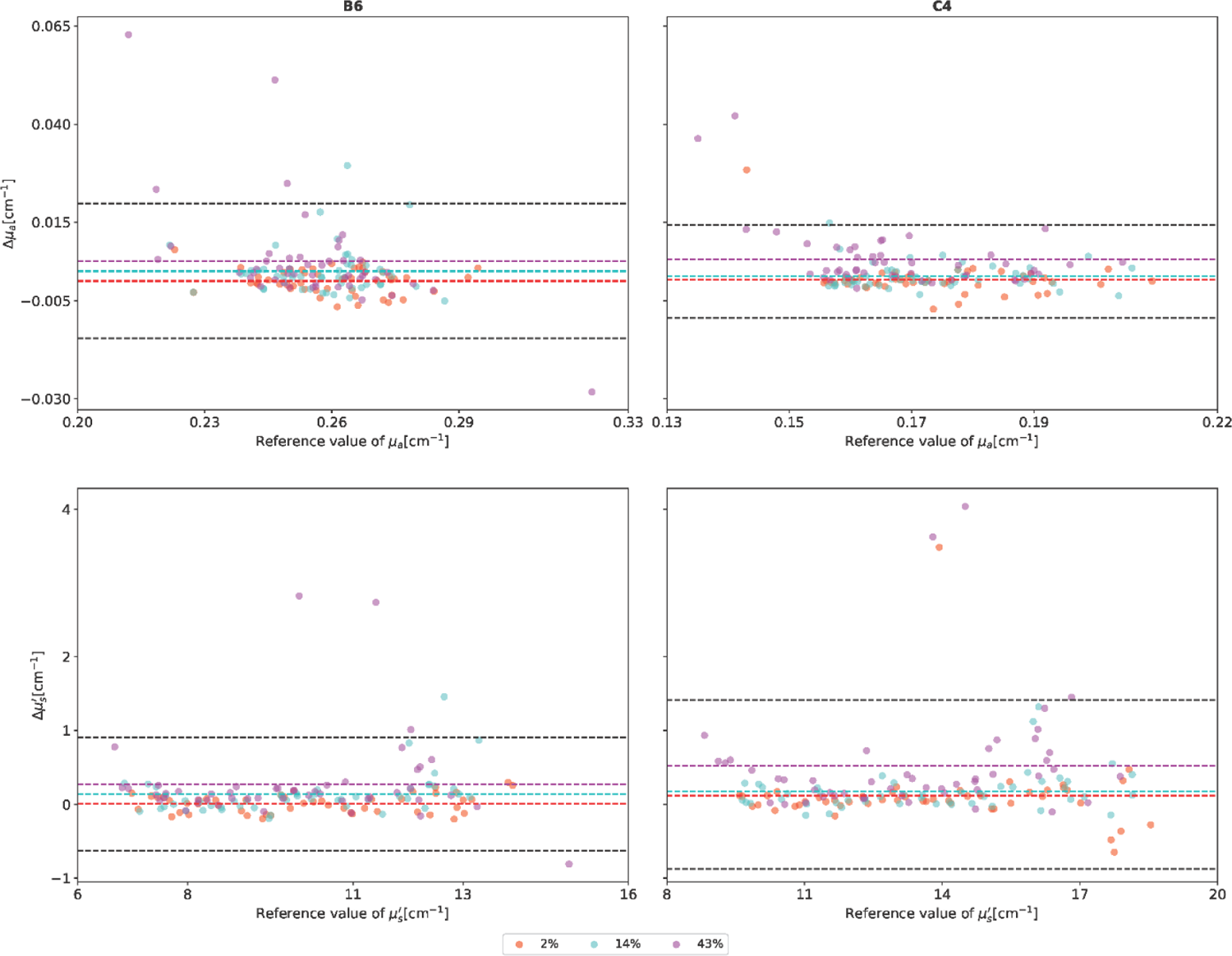
Bland Altman plots for the absorption coefficient (top row) and the reduced scattering coefficient (bottom row) spectra of phantom B6 (left column) and C4 (right column) for the skin phantoms mimicking different skin pigmentation tones (M_f_ = 0%, 2%, 14%, 43%). Colored horizontal dashed lines represent the bias (the average of the difference with the M_f_=0% case). On phantom B6, for μ_a_ (μ_s_’) the bias is 1.77 10^-5^ (0.0092), 0.0025 (0.13), and 0.0052 (0.27) cm^-1^ respectively for M_f_ = 2%, 14%, and 43%. On phantom C4, for μ_a_ (μ_s_’) the bias is 0.00040 (0.12), 0.0012 (0.17), and 0.0056 (0.52) cm^-1^ respectively for M_f_ = 2%, 14%, and 43%. Black horizontal dashed lines represent the upper and lower 95% limits of agreement (bias ± 1.96 × standard deviation of the difference).

For the case of M_f_ = 43% the bias (i.e., the mean value of the difference between the measured parameter obtained when placing a given skin phantom under the probe and the corresponding value when the control skin phantom with M_f_ = 0% is used) for phantom B6 (C4) is 0.0052 (0.0056) cm^-1^ and 0.27 (0.52) cm^-1^ for μ_a_ and μ_s_’, respectively. Much lower biases are obtained for the other M_f_ values. Apart from a few outliers, the difference in values is always within the 95% limits of agreement (i.e., bias ± 1.96 × standard deviation) and does not show any particular trend.

Besides being used to mimic static optical properties, bulk phantoms B6 and C4 can also be used to mimic static hemodynamic properties by assuming that the measured absorption coefficients at 686 nm and 830 nm of the phantoms are determined by a linear combination of O_2_Hb and HHb. Table 1 shows the results of TD NIRS measurements on bulk phantoms B6 and C4 with different pairs of skin phantoms on top, always keeping the control skin phantom M_f_ = 0% as one of the pair.

**Table 1.**
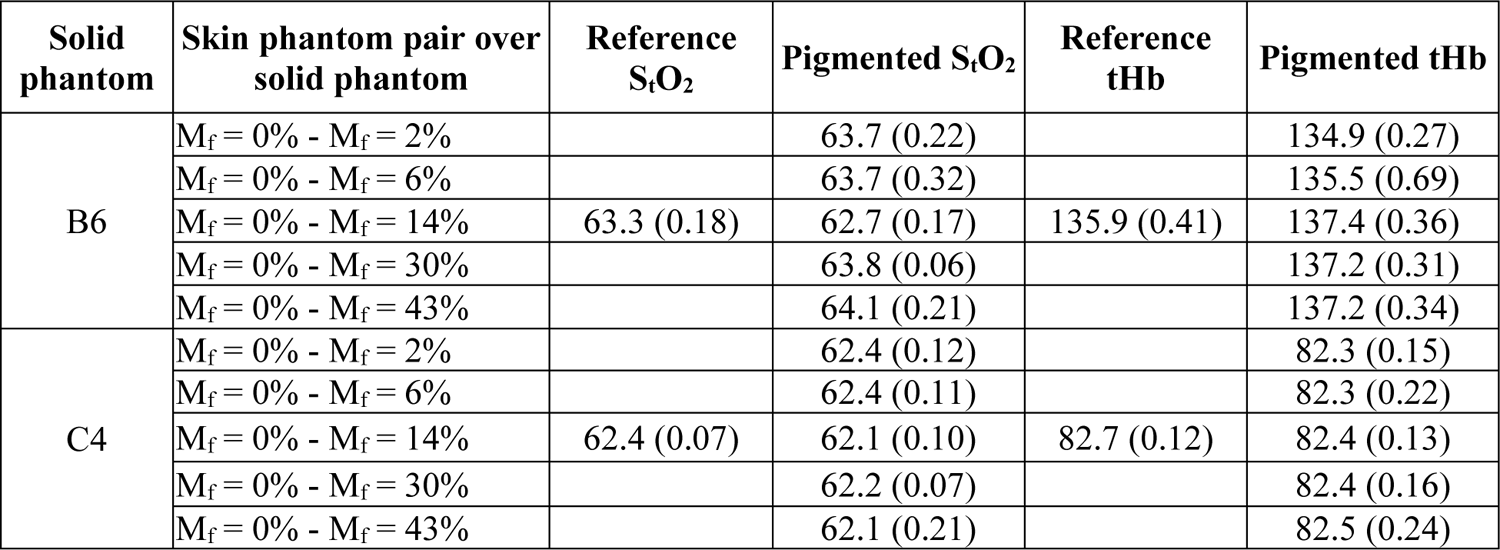
Estimated hemodynamic parameters, measured with the NIRSBOX oximeter superposing skin phantoms on B6 and C4 bulk phantoms: for the pigmented values we report the mean (standard deviation) over four different phantoms replicas while for the reference measurement we report the mean (standard deviation) over repositioning across all measurements.

| Solid phantom | Skin phantom pair over solid phantom | Reference $\text{S}_t\text{O}_2$ | Pigmented $\text{S}_t\text{O}_2$ | Reference tHb | Pigmented tHb |
| --- | --- | --- | --- | --- | --- |
| B6 | $M_f = 0\% - M_f = 2\%$ | | 63.7 (0.22) | | 134.9 (0.27) |
| | $M_f = 0\% - M_f = 6\%$ | | 63.7 (0.32) | | 135.5 (0.69) |
| | $M_f = 0\% - M_f = 14\%$ | 63.3 (0.18) | 62.7 (0.17) | 135.9 (0.41) | 137.4 (0.36) |
| | $M_f = 0\% - M_f = 30\%$ | | 63.8 (0.06) | | 137.2 (0.31) |
| | $M_f = 0\% - M_f = 43\%$ | | 64.1 (0.21) | | 137.2 (0.34) |
| C4 | $M_f = 0\% - M_f = 2\%$ | | 62.4 (0.12) | | 82.3 (0.15) |
| | $M_f = 0\% - M_f = 6\%$ | | 62.4 (0.11) | | 82.3 (0.22) |
| | $M_f = 0\% - M_f = 14\%$ | 62.4 (0.07) | 62.1 (0.10) | 82.7 (0.12) | 82.4 (0.13) |
| | $M_f = 0\% - M_f = 30\%$ | | 62.2 (0.07) | | 82.4 (0.16) |
| | $M_f = 0\% - M_f = 43\%$ | | 62.1 (0.21) | | 82.5 (0.24) |

The reference S_t_O_2_ and tHb values (estimated as average over all measurements with the control skin phantom) are 63.3±0.18 % and 135.9±0.41 μM for phantom B6 (corresponding to coefficients of variation of 0.3% and 1% for S_t_O_2_ and tHb, respectively), and 62.4±0.07 % and 82.7±0.12 μM for phantom C4 (corresponding to coefficients of variation of 0.1% and 0.2% for S_t_O_2_ and tHb, respectively). When the pigmented skin phantoms are used, negligible changes in the estimated S_t_O_2_ and tHb values are observed. Average S_t_O_2_ values range from 62.7 to 64.1% in phantom B6 (coefficient of variation 0.8%) and from 62.1 to 62.4% in phantom C4 (coefficient of variation 0.2%). Similarly, average tHb values range from 134.9 to 137.4 μM for phantom B6 (coefficient of variation 0.8%) and from 82.3 to 82.5 μM in phantom C4 (coefficient of variation 0.1%).

A two-tails, paired t-test was performed comparing the reference values versus the pigmented ones across the pigmentations in the S_t_O_2_ and tHb groups over the two bulk phantoms (C4 and B6). Results showed p-values > 0.05 for all comparisons, showing no significant difference between reference and pigmented measured values.

### *In vivo* broadband TD NIRS static measurements on volunteers with skin phantoms

Figure 3 shows the absorption and reduced scattering spectra of subject S05 on forearm, forehead and abdomen with skin phantoms mimicking different skin pigmentation tones (M_f_ = 0%, 2%, 14%, 43%). Supplementary Figure 1 and 2 report the spectra for subjects S01−S06. In the μ_a_ spectra of the forearm the contribution of hemoglobin^35^ (with the decrease from 600 nm to 700 nm, the peak of HHb around 760 nm, and the broad O_2_Hb rise over 800 nm), lipid^36^ (with the characteristic peaks around 930 nm and 1050 nm) and water^37^ (with the broad peak around 980 nm) are clearly visible. In the forehead the lipid contribution is not observable, while it dominates in the case of the abdomen, as expected. The μ_s_’ spectra of all body districts show the characteristic decrease with the wavelength, with lower values for the forearm, due to the muscle fiber structure^3^. The effect of increased skin pigmentation (as determined by the skin phantoms) on the absorption values is minimal for all tissues in the region below 900 nm, with a limited effect around 980 nm for the forearm and the forehead. An effect can be appreciated in the scattering spectra where small homothetic variations are induced by the different skin pigmentation levels.

**Fig. 3.**
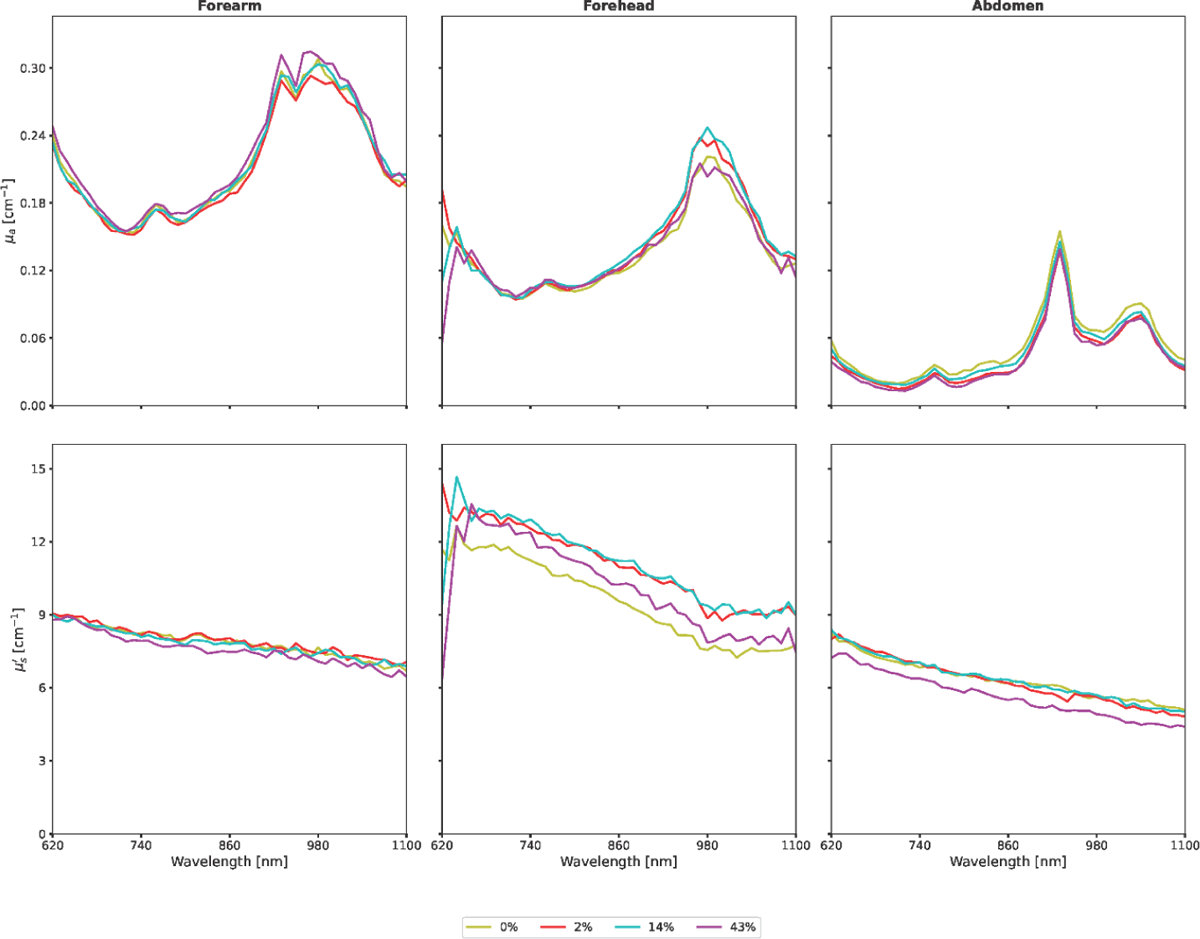
Absorption coefficient (top row) and reduced scattering coefficient (bottom row) spectra of subject S05 on forearm (left column), forehead (middle column), and abdomen (right column) with skin phantoms mimicking different skin pigmentation tones (M_f_ = 0%, 2%, 14%, 43%).

To better quantify the effect of the skin pigmentation, Figure 4 shows the Bland-Altman plots for μ_a_ and μ_s_’ for subject S05. The bias for μ_a_ in the forearm ranges from −0.0036 to 0.0021 cm^-1^ with the maximum values obtained at the lowest pigmentation level (M_f_ = 2%). In the forehead, the bias for μ_a_ ranges from −0.0040 to 0.0005 cm^-1^, while for the abdomen it spans from 0.0026 to 0.0056 cm^-1^. In the case of the abdomen, we observe a difference in values slightly lower than the limit of agreements for the M_f_ = 43%. For the μ_s_’, we have larger biases especially in the forehead and in the abdomen with values in the range −0.77 to 0.34 cm^-1^.

**Fig. 4.**
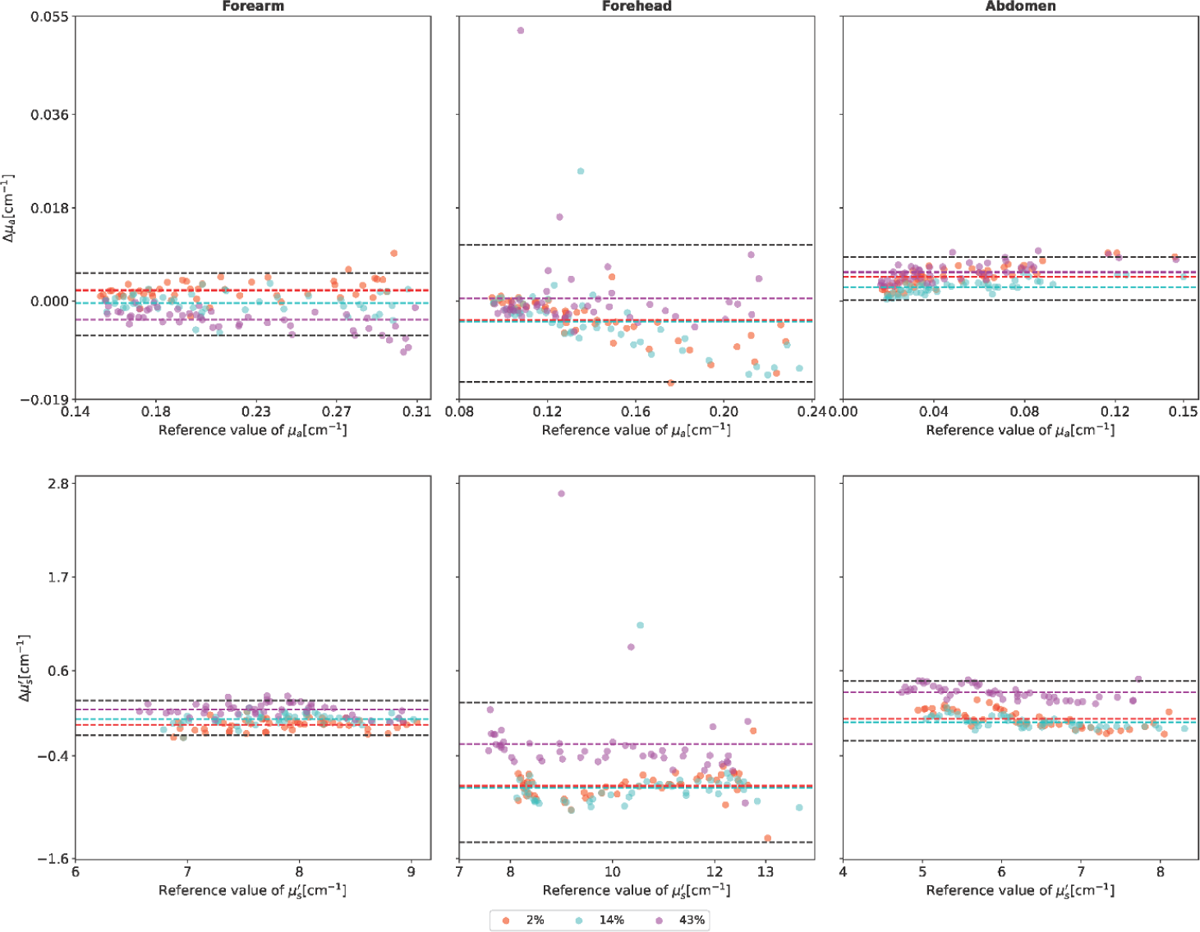
Bland-Altman plot for the absorption coefficient (top row) and the reduced scattering coefficient (bottom row) of subject S05 on forearm (left column), forehead (middle column), and abdomen (right column) with skin phantoms mimicking different skin pigmentation tones (M_f_ = 2%, 14%, 43%). Colored horizontal dashed lines represent the bias (the average of the difference with the M_f_ = 0% case). Black horizontal dashed lines represent the upper and lower 95% limits of agreement (bias ± 1.96 × standard deviation of the difference). On the forearm, for μ_a_ (μ_s_’) the bias is 0.0021 (−0.036), −0.0004 (0.031), and −0.0036 (0.15) cm^-1^ respectively for M_f_ = 2%, 14%, and 43%. On the forehead, for μ_a_ (μ_s_’) the bias is −0.0037 (−0.75), −0.0040 (−0.77), and 0.0005 (−0.76) cm^-1^ respectively for M_f_ = 2%, 14%, and 43%. On the abdomen, for μ_a_ (μ_s_’) the bias is 0.0047 (0.037), 0.0026 (−0.0042), and 0.0056 (0.34) cm^-1^ respectively for M_f_ = 2%, 14%, and 43%.

Figure 5 shows the corresponding Bland-Altman plots for subjects S01−S06. The bias for μ_a_ in the forearm ranges from −0.0075 to −0.0044 cm^-1^ with the minimum values obtained at the lowest pigmentation level (M_f_ = 2%). In the forehead the bias for μ_a_ ranges from 0.0031 to 0.0071 cm^-1^, while for the abdomen it spans from −0.00060 to −0.0022 cm^-1^. Like in the phantom study, the difference in values is mostly confined within the limits of agreement with no appreciable trend. For the μ_s_’ we have a larger bias with maximum values of 0.38 cm^-1^, 0.26 cm^-1^, and 0.11 cm^-1^ in the forearm, forehead and abdomen, respectively. A peculiar behavior is observed at M_f_ = 2% in the forearm and the abdomen with clustered values of the difference.

**Fig. 5.**
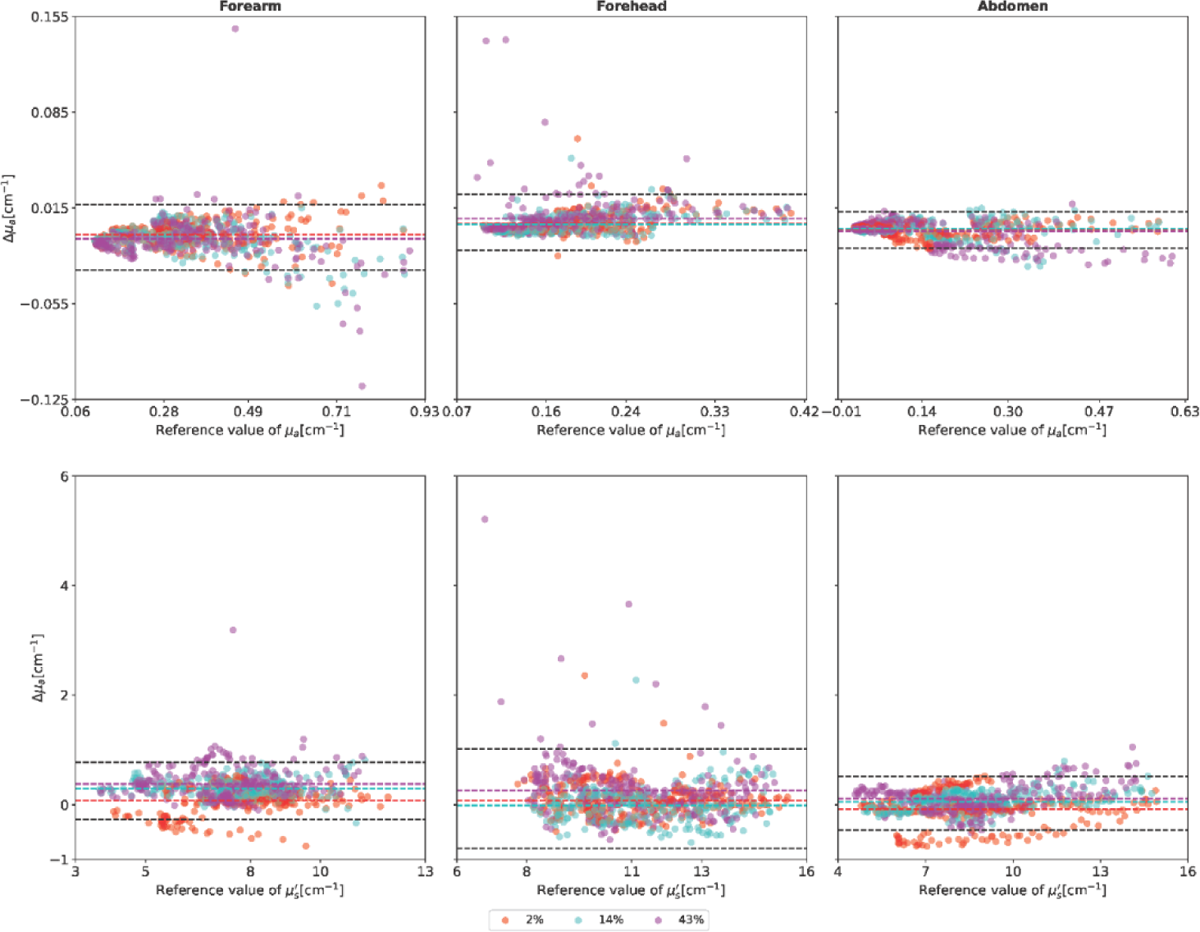
Bland-Altman plot for the absorption coefficient (top row) and the reduced scattering coefficient (bottom row) of subjects S01-S06 on forearm (left column), forehead (middle column), and abdomen (right column) with skin phantoms mimicking different skin pigmentation tone (M_f_ = 2%, 14%, 43%). Colored horizontal dashed lines represent the bias (the average of the difference with the M_f_ = 0% case). Black horizontal dashed lines represent the upper and lower limits of agreement (bias ± 1.96 × standard deviation of the difference). On the forearm, for μ_a_ (μ_s_’) the bias is −0.0044 (0.072), −0.0075 (−0.29), and −0.0075 (0.38) cm^-1^ respectively for M_f_ = 2%, 14%, and 43%. On the forehead, for μ_a_ (μ_s_’) the bias is 0.0035 (0.074), 0.0031 (−0.012), and 0.0071 (0.26) cm^-1^ respectively for M_f_ = 2%, 14%, and 43%. On the abdomen, for μ_a_ (μ_s_’) the bias is −0.0006 (−0.083), −0.0003 (0.052), and −0.0022 (0.11) cm^-1^ respectively for M_f_ = 2%, 14%, and 43%.

### Clinical TD NIRS static measurements on pediatric population

To evaluate the TD NIRS technique on a wider set of subjects, with a true representation of all skin pigmentation tones, clinical TD NIRS oximetry measurements were performed on 352 pediatric subjects in rest conditions without any pathological evidence. Clustering the entire cohort into different phototypes groups (using the Fitzpatrick scale as assessed by the clinicians) resulted in six distinct groups with varying population sizes: 17, 111, 129, 62, 26, and 7 subjects, corresponding to Fitzpatrick scores 1 through 6, respectively. Figure 6 shows S_t_O_2_ and tHb values obtained by TD NIRS in the forehead and in the middle upper arm as a function of the Fitzpatrick scores.

**Fig. 6.**
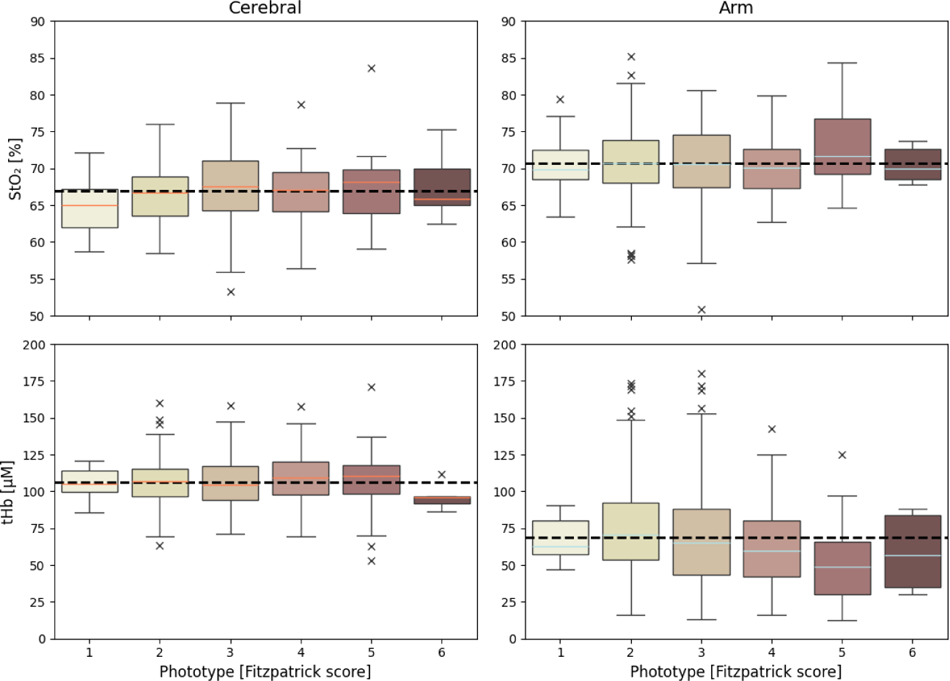
S_t_O_2_ (upper row) and tHb (lower row) measurements in the left frontotemporal region (left side) and middle upper arm (right side) of 352 pediatric subjects grouped according to the Fitzpatrick score. The central line within each box represents the median value, indicating the 50th percentile of the data. The lower and upper boundaries of the boxes correspond to the 25th (Q1) and 75th (Q3) percentiles, respectively, defining the interquartile range (IQR). Whiskers extend from the box edges to the smallest and largest observations within 1.5 times the IQR from Q1 and Q3, respectively. Data points beyond the whiskers are depicted as individual markers (“x”) and are considered outliers, indicating values that significantly deviate from the central distribution.

For both S_t_O_2_ and tHb the average values (dashed lines) are within the standard deviations of all groups. One-way ANOVA shows that there are no significant differences (p-value > 0.05) among the different Fitzpatrick group mean values in both S_t_O_2_ and tHb (arm S_t_O_2_: f-value = 1.10, p-value = 0.36; arm tHb: f-value = 1.13, p-value = 0.09; forehead S_t_O_2_: f-value = 1.24, p-value = 0.29; forehead tHb: f-value = 0.79, p-value = 0.56).

### *In vivo* TD NIRS dynamic measurements during arterial occlusion on volunteers with skin phantoms

In the previous sections we have shown the effect of skin pigmentation on TD NIRS measurements on bulk phantoms and *in vivo* in static conditions, i.e., with fixed optical and hemodynamic properties. In this section we illustrate the case of dynamic changes in the hemodynamic properties to emulate a possible pathological condition. For each subject, two measurements were performed synchronously, using two skin phantoms (control and pigmented) positioned adjacent to each other on the forearm.

Figure 7 reports the time evolution of S_t_O_2_ and tHb and the corresponding Bland-Altman plots for subject S11 during an arterial occlusion achieved through the inflation of a cuff and performed with different pairs of skin phantoms placed between the TD NIRS probe and the tissue. Supplementary Figure 3−7 report the same type of plots for all the other subjects (S07−S10 and S12).

**Fig. 7.**
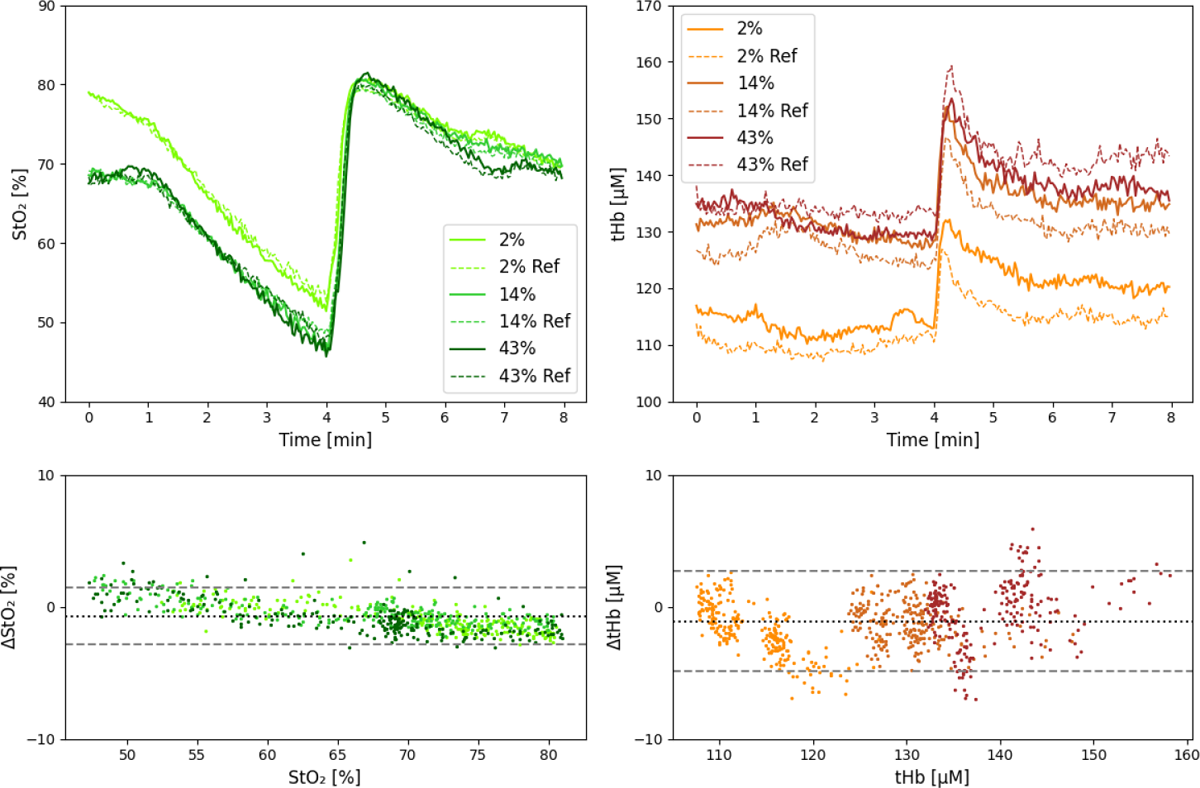
S_t_O_2_ (top left) and tHb (top right) during arterial occlusion in subject S11 with different skin phantoms (M_f_ = 2%, 14% and 43%, solid lines) against the simultaneous reference measurement (Ref, dashed lines). Bland-Altman plots for S_t_O_2_ (bottom left) and tHb (bottom right) are also reported taking into consideration all the datapoints of the arterial occlusion protocol. Black horizontal dashed lines represent the upper and lower 95% limits of agreement (bias ± 1.96 × standard deviation of the difference).

As expected, S_t_O_2_ decreases during the arterial occlusion and rapidly increases when the occlusion is removed. The difference in the baseline data for the case M_f_ = 2% with respect to the cases M_f_ = 14% and 43% is likely due to the different hemodynamic conditions of the arm since the three measurements were made in series (with some time interval in between). This can also be confirmed by the temporal evolution of tHb with reduced tHb values for the M_f_ = 2% case. The effect of the arterial occlusion on tHb, with a rather constant trend during cuff inflation and a rapid increase following cuff release, is expected. Differently from what happens for S_t_O_2_, here we notice a larger difference between control and pigmented measurements.

The absolute S_t_O_2_ bias between the measurement with the pigmented skin phantoms and the control skin phantom is <1%, indicating a small systematic difference between the configurations. The limits of agreement are relatively small as compared to the scale of the measured values, indicating a good numerical agreement between reference and pigmented measurements. We notice a trend for the S_t_O_2_ difference as function of the mean S_t_O_2_, indicating a small overestimation for S_t_O_2_ values below 70%, and underestimation above 70%. This deviation seems consistent across different pigmentations; therefore, it might be attributed to different physiological behaviors in the two nearby forearm locations. In the Bland-Altman plot for tHb we noticed that data for the different pigmentations are clustered around different mean values, however mostly confined within the limits of agreement.

In Figure 8 the Bland-Altman plots for S_t_O_2_ and tHb of all subjects during the full duration of the dynamic test are reported. For tHb measurements, physiological variability and probe repositioning appear to play a more significant role than pigmentation variations, making the contribution of different pigmentation levels negligible in static and dynamic tHb measurements. For S_t_O_2_, remarkably similar trends are observed between the pigmented and reference locations during the occlusion protocol across all subjects, despite diverse hemodynamic behaviors seen in consecutive measurements. The Bland-Altman plot shows that the deviation at the pigmented location is almost always within the limits of agreement throughout the occlusion protocol. The only instances where the limits of agreement are exceeded occur during the reoxygenation process, characterized by high S_t_O_2_ variations within a noticeably short time interval. This rapid hemodynamic process (less than 3 s) is comparable to the acquisition frequency (1 Hz). The low acquisition rate and possible physiological differences between the two measurement locations can explain the observed outliers in the Bland-Altman plots.

**Fig 8.**
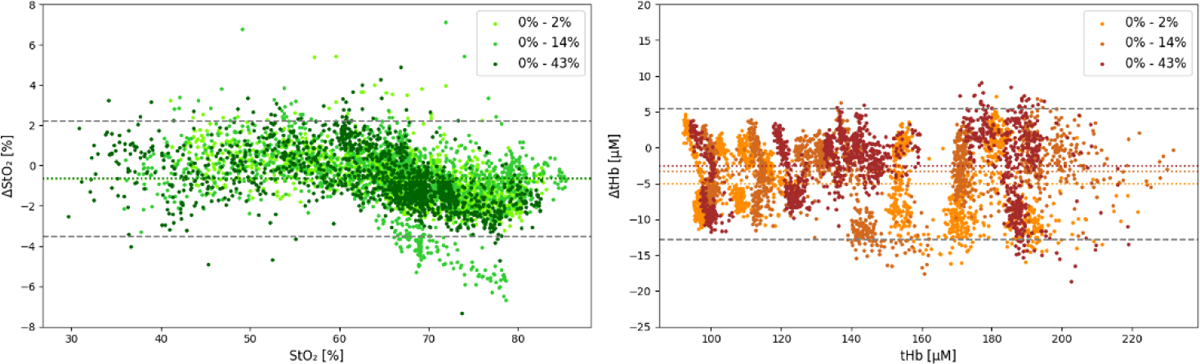
Bland-Altman plot for S_t_O_2_ (left) and tHb (right) of subject S07-S12 during arterial occlusion with skin phantoms mimicking different skin pigmentation tone (M_f_ = 2%, 14%, 43%). Colored horizontal dashed lines represent the bias (the average of the difference with the M_f_ = 0% case). Black horizontal dashed lines represent the upper and lower 95% limits of agreement (bias ± 1.96 × standard deviation of the difference).

To further quantify the possible effect of skin pigmentation during the vascular occlusion test, we report in Table 2 the average changes (and the corresponding standard deviation) across all subjects S07−S12 for parameters that are typically estimated in a vascular occlusion test: desaturation slope, reoxygenation slope, and area under the curve (AUC) in the recovery phase. The change in the desaturation slope seems minimally affected by pigmentation, being −0.5 %/min for M_f_ = 2% and 14 % and −0.6 %/min for M_f_ = 43%. The reoxygenation slope has a clearer trend with pigmentation being −3.6%/min for Mf = 2% and increasing to 3.2 %/min and 4.5 %/min for M_f_ = 14% and 43%, respectively. The change in AUC ranges from 1.1 to 1.9 % min with no correlation with increasing pigmentation. However, results of the statistical ANOVA test show that no significant effect to the figures of merit in the occlusion protocol can be referred to different pigmentation levels of the skin phantoms.

**Table 2.** Average changes and standard deviation across subjects S06-S12 for desaturation slope, reoxygenation slope, and area under the curve (AUC). Results of the statistical ANOVA test are also reported.

| Skin phantom pair over the arm | $\Delta$ Desaturation Slope [%/min] | $\Delta$ Reoxygenation Slope [%/min] | $\Delta$ AUC [% x min] |
| --- | --- | --- | --- |
| $M_f = 0\% - M_f = 2\%$ | -0.5 (0.2) | -3.6 (11.2) | 1.9 (1.3) |
| $M_f = 0\% - M_f = 14\%$ | -0.5 (0.9) | 3.2 (10.2) | 1.1 (1.0) |
| $M_f = 0\% - M_f = 43\%$ | -0.6 (0.9) | 4.5 (12.8) | 1.4 (0.6) |
| ANOVA *p-value* | 0.97 | 0.49 | 0.42 |
| ANOVA *f-value* | 0.03 | 0.72 | 0.93 |

## Discussion

We investigated the effect of skin pigmentation on TD NIRS measurements. This study holds significant appeal for the wider biomedical community: there is in fact a growing interest in evaluating the effects of skin pigmentation on light-based techniques as demonstrated by recent publications on pulse oximetry^38^. While targeting different physiological parameters, pulse oximetry and NIRS indeed share the same physical principles regarding light propagation in biological tissues, as they both rely on illuminating the tissue with NIR light and detecting the reemitted (backscattered or transmitted) photons.

The approach we followed involved several steps: i) the fabrication of tissue-mimicking skin phantoms with variable concentration of nigrosine to mimic the absorption by melanin; ii) the use of skin phantoms for static TD NIRS measurements on calibrated bulk phantoms mimicking tissue optical properties; iii) the use of skin phantoms for *in vivo* TD NIRS measurements on volunteers during static (at rest) measurements; iv) the use of skin phantoms for *in vivo* TD NIRS measurements on volunteers during dynamic (arterial occlusion) protocols.

The fabrication and use of skin-mimicking phantoms (and of phantoms in general) for testing basic performance of optical technologies must be preferred to the sole *in vivo* testing, thanks to reduced complexity and risks and to the possibility to facilitate quality assurance measurements and device comparison for intra and inter laboratory measurements. The disadvantage of phantoms is related to the scarce availability of phantom producers on the market that forces researchers to develop the required expertise in-house. Thanks to our long-term expertise in phantom preparation^39-41^, we were successful in realizing skin phantoms for NIRS measurements. Furthermore, we have utilized the valuable results of other research groups^26-28^, in particular for the choice of using nigrosine to mimic skin pigmentation and for the reference optical properties of the skin phantoms.

We have first studied the effect of pigmentation in static conditions by broadband TD NIRS measurements. Apart from the intrinsic simplicity in performing static (versus dynamic) measurements, the motivation behind this approach is that diffuse light is also used in clinical applications like optical mammography, where minimal perturbation of the optical properties is expected, and the overall performances of the technique are mostly determined by the ability to assess static optical properties. From these measurements we have observed that the effect of skin pigmentation on TD NIRS-derived parameters is negligible for μ_a_ in the spectral region below 900 nm, while care should be taken for longer wavelengths especially at the highest pigmentation level since the combination of a naturally occurring absorption (due to water and lipid) may add to the absorption of melanin. The combined effect is a strong reduction of the TD NIRS signal-to-noise-ratio (SNR). For static measurements, the SNR reduction could be partially overcome by increasing the acquisition time, if the background noise is relatively low. A larger effect of skin pigmentation is observed on μ_s_’, but the amount is relatively small. The larger absolute difference is <1 cm^-1^ resulting in a relative difference <20% in the worst case of μ_s_’ = 5 cm^-1^. Anyway, small changes in the scattering coefficient have limited influence on the characteristics of light propagation in diffusive media, but they might affect the estimate of parameters derived by CW NIRS if not properly considered^42^.

Static measurements on bulk phantoms mimicking hemodynamic parameters in the arm and in the head were used to assess the influence of skin pigmentation on S_t_O_2_ and tHb as derived by TD NIRS oximetry. The estimate of S_t_O_2_ and tHb obtained by TD NIRS oximetry in static conditions are substantially unaffected by skin pigmentation. Similar results were obtained in the *in vivo* measurements performed in the clinical settings on pediatric patients.

Raw data and statistical tests showed that in dynamic measurements the different skin pigmentation tones have negligible effect on S_t_O_2_. This is perhaps the most valuable result of this study since it underscores the reliability and potential of TD NIRS in tissue oximetry. Differently from CW NIRS^28-30^ or photoacoustic imaging^26-27^, in TD NIRS no significant bias on S_t_O_2_ is introduced by skin pigmentation. Conversely, the higher variability of tHb results observed during the dynamic measurements can be explained not only by an incomplete hemodynamic recovery between successive measurements in the same subject, but also by the larger sensitivity to probe repositioning and the physiological condition of the subjects. Trends for tHb also differ between the pigmented region and the reference region during simultaneous acquisitions, regardless of M_f_ values. The Bland-Altman results did not show consistent behavior across different measurements, preventing any definitive conclusions for tHb. The reference (non-pigmented) measurement was performed in a different location, although near to the pigmented measurement. The difference position might lead to a slight difference in the hemodynamic behavior, due to physiological variability in the acquisition. It would be best to compensate for this physiological variability by increasing the number of subjects measured and randomizing the probe positioning of reference and pigmented measurements.

We acknowledge some limitations in this study. We have not used numerical simulations (e.g., by Monte Carlo method in complex geometries) since we believe that calibrated tissue phantoms can be a valid alternative (provided that the measuring setup is performant). Only one subject (S05=S11) underwent both static and dynamic TD NIRS measurements. Indeed, the aim was not comparing the performance of the two TD NIRS devices (the broadband TD NIRS system and the commercial TD NIRS tissue oximeter) on the same subject, rather we wanted to assess the effect of the skin pigmentation on a large variety of subjects. For the same reason, we have not considered selecting subjects based on anatomical or physiological characteristics (e.g., the thickness of the adipose tissue covering the muscle). Indeed, we have not objectively evaluated the skin pigmentation level in our subjects (e.g., by melanometry as suggested by Ref. 38). However, we notice that we have used the skin phantoms mimicking different pigmentation levels on top of the natural skin of the subjects, therefore potentially creating an even darker pigmentation as compared to natural pigmentation. Regarding the TD NIRS clinical campaign, a notable limitation is the uneven distribution of subjects across the different Fitzpatrick phototypes. The enrolled subjects were not homogeneously distributed among all pigmentation categories, which could potentially introduce bias and affect the generalizability of our findings. Despite the distributional imbalance, the results contribute meaningful information to the understanding of measurement biases in TD NIRS and their relation to skin pigmentation. Future studies should aim for a more balanced *in vivo* sample across all Fitzpatrick phototypes to enhance the robustness and applicability of the findings.

Overall, we reliably demonstrated through phantoms and *in vivo* measurements in static and dynamic conditions that skin pigmentation does not significantly affect the estimate of optical properties and tissue oximetry parameters by TD NIRS, underscoring the reliability and potential of this emerging technology in diverse clinical settings.

## Materials and methods

### Skin phantom preparation

Thin tissue-mimicking phantoms with different skin pigmentation ranging from light to dark (and including a control phantom with no pigmentation), corresponding to M_f_ = 0%, 2%, 6%, 14%, 30%, and 43%, were prepared by mixing silicone (Silicone Elastomer Sylgard 184, Base 10 and Curing Agent 1, Dow Corning) as matrix, TiO_2_ powder (T-8141, Sigma–Aldrich, St. Louis, Missouri) as scattering agent, and a solution of alcohol-soluble nigrosine (211680, Merck Life Science S.r.l., Italy) in ethanol (414605, Carlo Erba Reagents S.A.S., France) as absorber. Table 3 reports composition and nominal optical properties at 685 nm of the phantoms. The mixtures were degassed in vacuum and small amounts were poured into specially designed metal moldings for oven curing at 60 °C for 6 hours. Four replicas of each skin phantom were prepared. The average thickness of the phantoms (evaluated using a digital caliper, taking six measurements on each layer, radially spaced by an angle of 60°) was 270 ± 10 μm. The average diameter of the skin phantoms was 60 mm. Finally, the skin phantoms were also divided into two half disks as shown in Fig.9 to ease the realization of a pair of skin phantoms (see Supplementary Fig. 8).

**Fig. 9:**
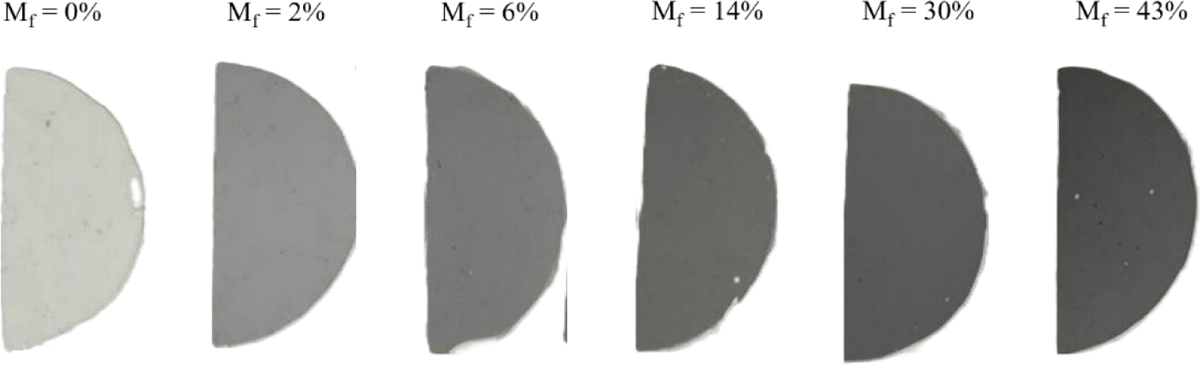
Picture of the skin phantoms.

**Table 3.** Composition and nominal optical properties of the skin pigmentation phantoms.

| $M_f$<br>[%] | Nigrosine<br>[mg] | Ethanol<br>[ml] | Silicone<br>base<br>[g] | Curing<br>agent<br>[g] | $TiO_2$<br>[mg] | nominal<br>$\mu_a$ [$cm^{-1}$] | nominal<br>$\mu_s'$ [$cm^{-1}$] |
| --- | --- | --- | --- | --- | --- | --- | --- |
| 0 | 0.00 | 0.7 | 32 | 3.2 | 59.8 | 0.0 | 20 |
| 2 | 1.95 |  |  |  |  | 3.4 |  |
| 6 | 5.87 |  |  |  |  | 10.2 |  |
| 14 | 13.77 |  |  |  |  | 23.9 |  |
| 30 | 29.53 |  |  |  |  | 51.2 |  |
| 43 | 42.33 |  |  |  |  | 73.5 |  |

### Broadband TD NIRS device (PoliMi)

The laboratory-grade broadband TD NIRS device is based on a supercontinuum fiber laser (SuperK Extreme FIR-20, NKT Photonics, Denmark) emitting pulses at a repetition rate of 40 MHz in the wavelength range 550-1750 nm. Wavelength selection is achieved through the rotation of a Pellin-Broca prism; afterwards, light is coupled to an optical fiber, attenuated by a neutral density filter, and delivered to the sample. Diffuse light coming from the sample is collected in reflectance geometry through a second fiber (at a distance of 2.5 cm from the injection fiber) which sends it to the detector, a Silicon Photo-Multiplier (SiPM, S10362-11-050C, Hamamatsu). DTOFs are reconstructed by a time-correlated single-photon counting board (SPC-130, Becker & Hickl, Germany). Further details on the setup can be found in literature^43-45^.

### TD NIRS tissue oximeter (NIRSBOX, PIONIRS Srl)

A commercially available, research-grade, TD NIRS tissue oximeter, (NIRSBOX, PIONIRS s.r.l., Italy) was also used^46^. The device uses two picosecond diode lasers emitting at 686 nm and 830 nm, a single-photon detector (SiPM, with optical filters to reduce ambient light noise), and timing electronics (time-to-digital converter, with 9.7 ps/ch resolution) to record the DTOF for the photons re-emitted from the tissue. The Instrument Response Function (IRF) has a full-width-at-half-maximum (FWHM) of about 150 ps. Two identical NIRSBOX devices (S/N: 009 and S/N: 015) were used^47^. The laser pulses emitted by the two devices were interleaved to avoid any possible cross-talk between the two devices. A custom probe hosting two pairs of source-detector optical fibers was employed. The optical probe was designed setting a distance of 2 cm between the source-detector pairs for the different NIRSBOX devices to maintain close proximity between the two measurement locations.

### TD NIRS static measurements with skin phantoms on bulk phantoms

Two sets of TD NIRS measurements were performed with the skin phantoms sequentially positioned on top of two solid bulk phantoms (labelled B6 and C4) mimicking the bulk optical properties of biological tissue^35^. The bulk phantoms B6 and C4 are made of epoxy resin (NM500/H179B; Nils Malmgren AB, Ytterby, Sweden) as matrix, TiO_2_ powder (T-8141, Sigma–Aldrich, St. Louis, Missouri) as scatterer, and black toner powder (black 46/I, part No. 88598306, Infotec, France) as absorber. They are cylinders (4.5 cm height and 10.5 cm diameter) with nominal optical properties (μ_a_ and μ_s_’) at 800 nm of 0.25 cm^-1^ and 10 cm^-1^ for phantom B6, and 0.15 cm^-1^ and 15 cm^-1^ for phantom C4.

First, broadband TD NIRS measurements were performed in the wavelength range 620-1100 nm, in steps of 10 nm with an overall acquisition time of 1 s per wavelength, after placing skin phantoms M_f_ = 0%, 2%, 14% and 43% on top of bulk phantoms B6 and C4. Then, TD NIRS measurements at 686 nm and 830 nm with 1 s acquisition time were performed using two identical TD NIRS tissue oximeters. A first oximeter performed measurements after sequentially placing skin phantoms M_f_ = 2%, 14%, 30% and 43% on top of bulk phantoms B6 and C4. In parallel, a second oximeter performed TD NIRS measurements on the control skin phantom (M_f_ = 0%) positioned on the same bulk phantom near to the other skin phantom (see Supplementary Figure 8 for details). The measurements with the TD NIRS tissue oximeters were repeated for the four different replicas of the skin phantoms to estimate the dispersion of measured parameters due to probe replacement and skin phantom fabrication. Data acquired by the TD NIRS tissue oximeters were interpreted as if the measured samples were biological tissue, which is assuming O_2_Hb and HHb as unique chromophores contributing to absorption. Indeed, the nominal optical properties of bulk phantom B6 are similar to muscle tissue corresponding to equivalent values of S_t_O_2_ and tHb equal to 62% and 134 μM, respectively, while bulk phantom C4 is similar to head tissue with equivalent values of S_t_O_2_ and tHb equal to 62% and tHb of 83 μM, respectively.

### *In vivo* broadband TD NIRS static measurements on volunteers with skin phantoms

We measured a total of 6 healthy volunteers (see Table 4 for details on gender and skin pigmentation) in three different body locations: forearm, forehead, and abdomen. The degree of pigmentation of each subject was determined in terms of Monk pigmentation score through visual examination by direct comparison with a reference Monk chromatic scale^13^.

**Table 4.** Demographic and skin tone properties of the subjects measured by the broadband TD NIRS device.

| Subject ID | Gender | Age (y) | Monk Skin Tone |
| --- | --- | --- | --- |
| S01 | M | 27 | 3 |
| S02 | F | 39 | 3 |
| S03 | M | 29 | 3 |
| S04 | M | 26 | 3 |
| S05 | M | 26 | 6 |
| S06 | F | 25 | 7 |

For each location, broadband TD NIRS measurements were performed in reflectance geometry with a source-detector distance of 2.5 cm and in the wavelength range 620-1100 nm in 10 nm steps. The TD NIRS probe holding the injection and collection fibers was handheld by the subjects and the skin phantoms (M_f_ = 0%, 2%, 14%, and 43%) were interposed between the tissue and the TD NIRS probe.

### Clinical TD NIRS static measurements on pediatric population

We enrolled a cohort of pediatric participants, aged 0-18 years (see Table 5), admitted to the Pediatric Department of Buzzi Children’s Hospital in Milan, Italy, from March 2023 to February 2024. Candidates with stable clinical conditions have been considered encompassing the following inclusion criteria: absence of fever, absence of cardiac or pulmonary pathologies, no chronic diseases, no ongoing pharmacological treatments, stable vital parameters (heart rate, respiratory frequency and peripheral oxygen saturation, S_p_O_2_), absence of wounds in the measured position and confirmation of normal hematocrit levels through blood analyses. The study was conducted in accordance with the Helsinki Declaration of 1975, as revised in 2008. The institutional ethics committee approved the protocol (Ethics Committee Milano Area 1; Study Registration 2022/ST/229; Protocol No. 0004021/2023 Date 30/01/2023). After receiving information about the study, all participants or their guardians provided written consent.

**Table 5.** Descriptors of the pediatric population.

| Parameters | Mean | Std | Min | Max |
| --- | --- | --- | --- | --- |
| Age (years) | 8.4 | 5.0 | 0.1 | 17.9 |
| Weight (Kg) | 34.3 | 22.8 | 3.9 | 128.1 |
| Height (cm) | 127.1 | 30.7 | 49.0 | 187.0 |
| Mid-upper arm circumference (cm) | 20.7 | 5.4 | 11.0 | 42.0 |
| Head circumference (cm) | 52.2 | 3.9 | 36.0 | 62.0 |
| Nasion –inion distance (cm) | 36.0 | 4.1 | 19.0 | 46.0 |
| Periauricular points distance (cm) | 32.5 | 3.2 | 17.0 | 40.0 |

TD NIRS measurements were performed on two locations using the NIRSBOX oximeter, in combination with a G5 “Goccia” optical probe^48^, having an inter-fiber distance of 2.5 cm. Locations were: the left frontotemporal cortex (Fp1 of the 10/20 EEG system mapping) and the left mid-upper arm (below the deltoid region), targeting cerebral and peripheral muscle tissue hemodynamics, respectively.

### *In vivo* TD NIRS dynamic measurements during arterial occlusion on volunteers with skin phantoms

Six healthy volunteers (see Table 6) were recruited, and TD NIRS acquisitions were conducted on the first proximal third of the right forearm, medially, during a vascular occlusion test.

**Table 6.** Demographic and skin tone properties of the subjects measured by the TD NIRS tissue oximeter.

| Subject ID | Gender | Age (y) | Monk Skin Tone |
| --- | --- | --- | --- |
| S07 | M | 25 | 2 |
| S08 | M | 29 | 4 |
| S09 | M | 34 | 3 |
| S10 | F | 29 | 2 |
| S11 | M | 26 | 6 |
| S12 | F | 26 | 3 |

The experimental protocol had a total duration of 8 min, divided into three phases: i) Baseline (1 min): the volunteer remains at rest. ii) Arterial occlusion (3 min): a 250 mmHg occlusion performed with a manual pneumatic cuff is applied for 3 min to the right biceps. iii) Recovery (4 min): the cuff is released. During the acquisitions, the subject was asked to sit on a chair in a relaxed position with the forearm laying on a desk at the same height as the heart. The degree of pigmentation of each subject was determined in terms of Monk pigmentation score through visual examination by direct comparison with a reference Monk chromatic scale^13^. The setup consisted of two NIRSBOX tissue oximeters, a custom probe comprising two injection fibers and two detection fibers, set at a distance of 2.5 cm, and four skin phantoms representing M_f_ = 0%, 2%, 14%, and 43%. For each of the six subjects, three different acquisitions were performed, two skin phantoms (control and one of the pigmented) were positioned adjacent to each other on the forearm. The control skin phantom (M_f_ = 0%) was placed in the most proximal position, while the pigmented skin phantom (M_f_ = 2%, or 14%, or 43%) was placed in the distal location. The probe (i.e., injection and collection fibers) of the first TD NIRS oximeter was positioned over the control skin phantom, while the probe of the second TD NIRS oximeter was positioned over the pigmented phantom (see Supplementary Figure 8 for details). We chose to use just three pigmented skin phantoms corresponding to three M_f_ levels to avoid subjecting the volunteers to six different occlusions. A minimum of 15 min was allowed to pass between one measurement and the next, to allow for recovery. To ensure consistency in the placement of the skin phantoms between one measurement and the next, a make-up pencil was used to mark the outer edges. Moreover, the temporal order in which the different pigmented skin phantoms were placed on the volunteer’s forearm was randomized to avoid the results showing an influence linked to the number of consecutive occlusions performed.

For each subject, the following figures of merit are calculated: the desaturation slope, the reoxygenation slope and the AUC during the recovery phase. The desaturation slope is determined by linear fitting of the S_t_O_2_ curve from 30 s after the onset of occlusion until 30 s before the end of occlusion. The reoxygenation slope is assessed by linear fitting of the S_t_O_2_ curve between 10% and 90% of the maximum-to-minimum range of the reoxygenation curve. The hyperemic peak evaluation is performed by calculating the AUC where the hyperemic peak S_t_O_2_ curve exceeds the baseline value, as determined during the final period of recovery.

The difference between the figures of merit of the reference measurement vs the pigmentated one has been calculated for each subject and each pigmentation value. Averaged difference values and their standard deviations have been calculated across all subjects. To evaluate for variation due to pigmentation values, a one-way ANOVA test has been conducted comparing different pigmentation for each figure of merit.

### TD NIRS data analysis

Each raw DTOF acquired by both TD NIRS devices has been analyzed via the solution to the photon diffusion equation for a semi-infinite homogeneous medium with extrapolated boundary conditions, after convolution with the IRF to retrieve absolute values of μ_a_ and μ_s_’ of the tissue under investigation^49^. The best fit was obtained using a Levenberg-Marquardt algorithm by minimizing the cost function χ^2^ while varying both μ_a_ and μ_s_’. Fitting included all points in the DTOF between 90% of the leading edge and 1% of the trailing edge. When using the TD NIRS tissue oximeter, starting from μ_a_ at 686 nm and 830 nm, and by exploiting the Beer law^35^, the concentrations of HbO_2_ and HHb have been obtained and tHb and S_t_O_2_ calculated.

### Data availability

All the data and methods needed to evaluate the conclusions of this work are presented in the main text and Supplementary Materials. Additional data can be requested from the corresponding author upon reasonable request.

## Acknowledgements

We acknowledge the support of Giovanna Sgarzi during the preparation of the skin phantoms and during TD NIRS oximetry measurements.

## Author Contributions

ML: conceived and initiated the research, support in constructing the experimental set-up, TD NIRS oximetry measurements, manuscript writing, manuscript revision.

CA: TD NIRS oximetry data analysis, manuscript revision.

IB: broadband TD NIRS measurements, manuscript writing, manuscript revision. AB: broadband TD NIRS measurements, manuscript revision.

MB: conceived and initiated the research, support in constructing the experimental set-up, TD NIRS oximetry measurements, manuscript revision.

VC: clinical TD NIRS measurements, manuscript revision.

DC: conceived and initiated the research, TD NIRS oximetry data analysis, manuscript revision. VD: broadband TD NIRS measurements, manuscript revision.

FN: skin phantom preparation and characterization, manuscript revision. VR: clinical TD NIRS measurements, manuscript revision.

LS: TD NIRS oximetry data analysis, manuscript revision. SZ: clinical TD NIRS measurements, manuscript revision. GV: clinical TD NIRS measurements, manuscript revision.

AT: conceived and initiated the research, manuscript writing, manuscript revision.

## Supplementary information

### S1 Broadband TD NIRS spectra on volunteers

**Supplementary Figure 1:**
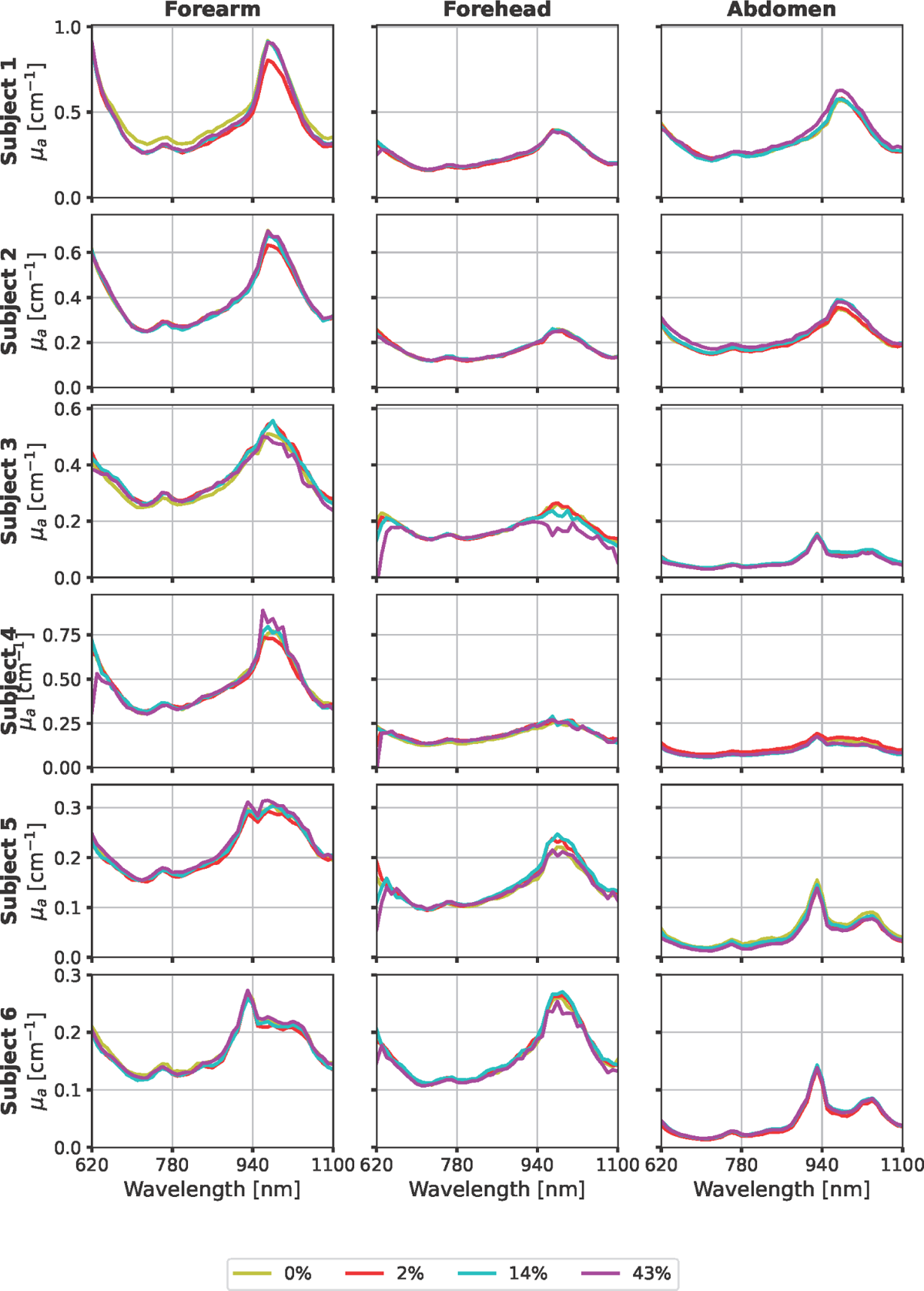
Absorption coefficient spectra on forearm (left column), forehead (middle column), and abdomen (right column) with skin phantoms mimicking different skin pigmentation tone (M_f_ = 0%, 2%, 14%, 43%). Each row corresponds to a different subject.

**Supplementary Figure 2:**
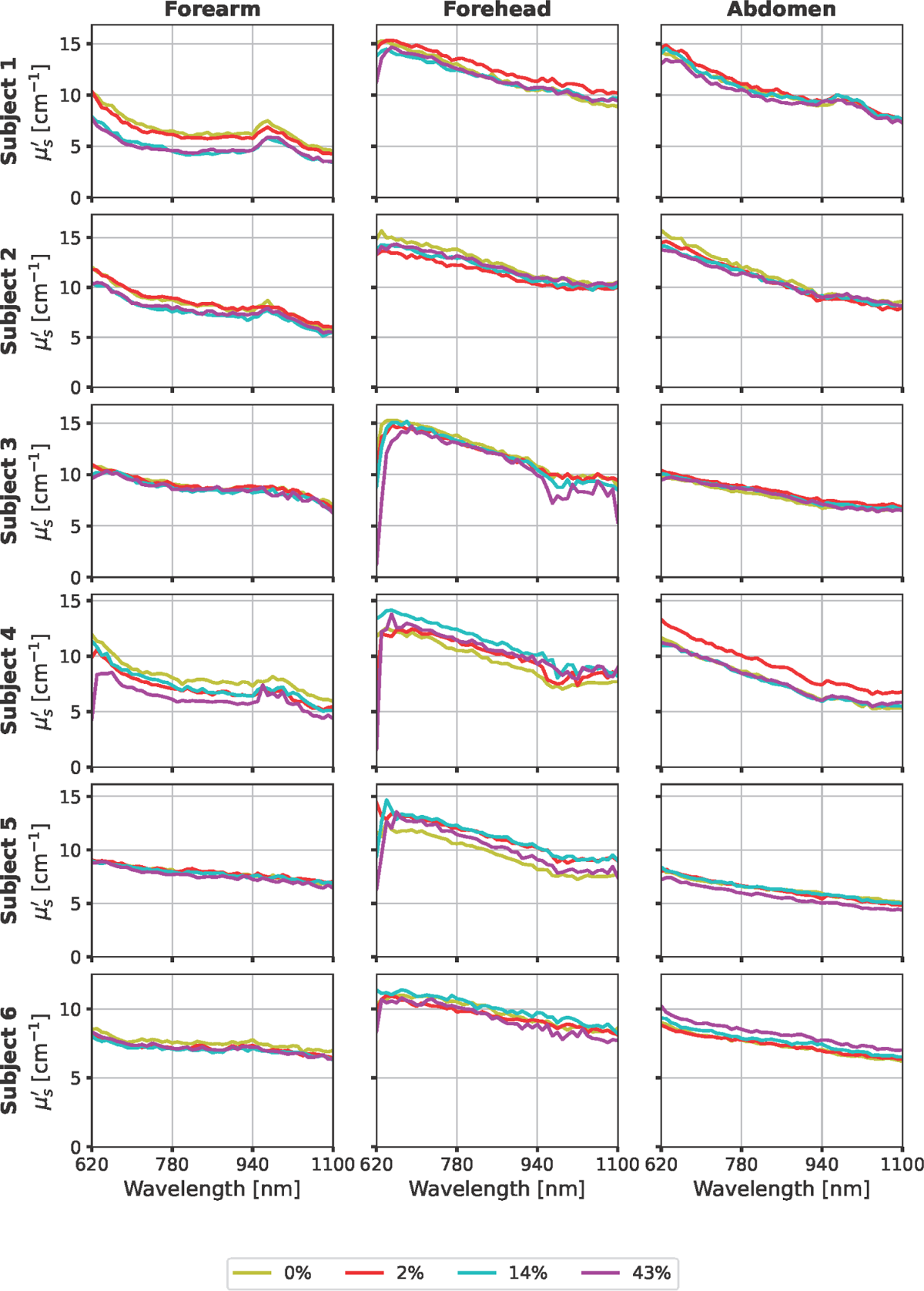
Reduced scattering coefficient spectra on abdomen (left column), forearm (middle column), and forehead (right column) with skin phantoms mimicking different skin pigmentation tone (M_f_ = 0%, 2%, 14%, 43%). Each row corresponds to a different subject.

### S2 TD NIRS arterial occlusion on volunteers

**Supplementary Figure 3:**
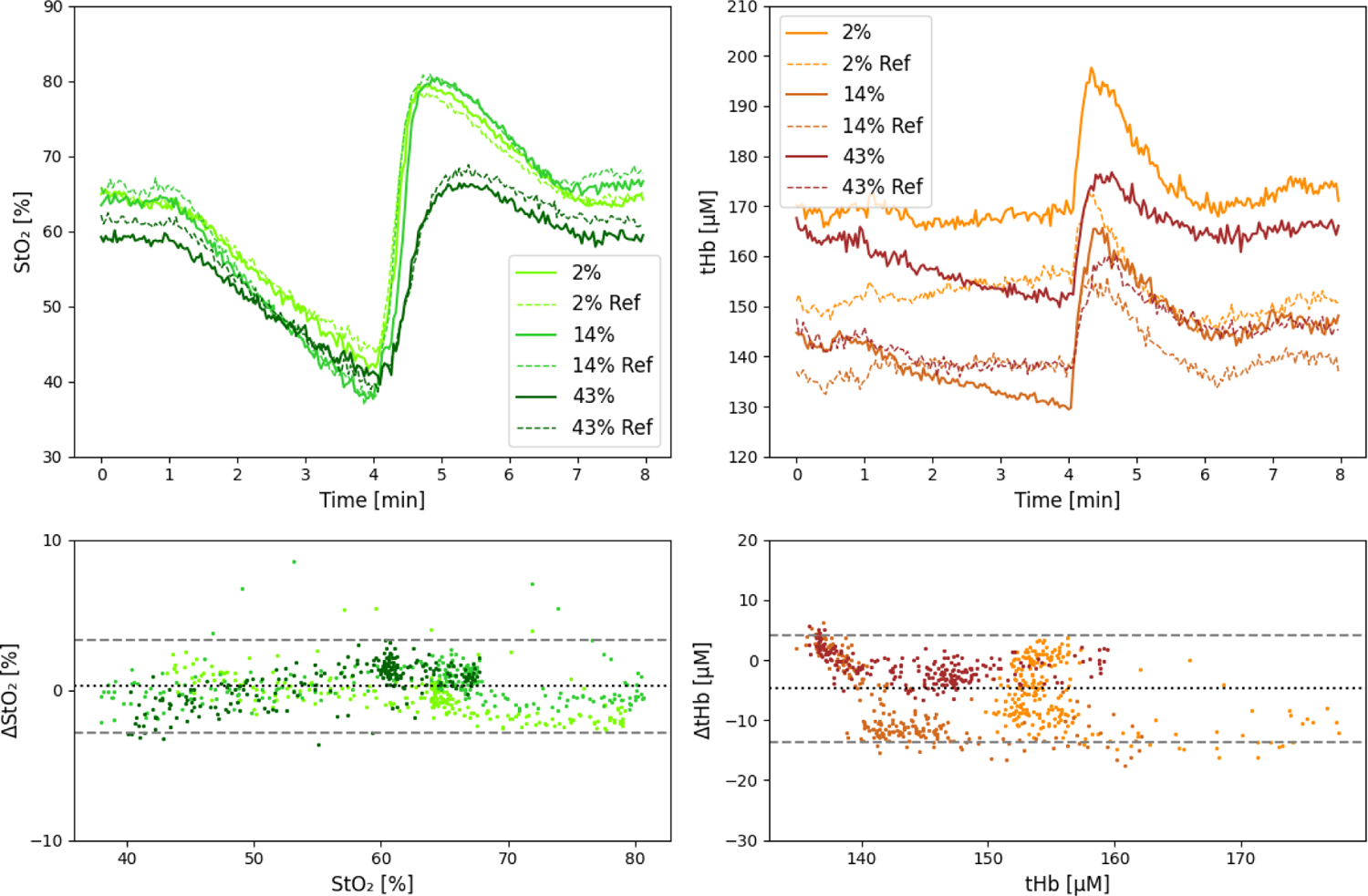
Time courses (top row) and Bland Altman plot (bottom row) for S_t_O_2_ (left column) and tHb (right column) during arterial occlusion with skin phantoms mimicking different skin pigmentation tone (M_f_ = 0%, 2%, 14%, 43%) for subject S07.

**Supplementary Figure 4:**
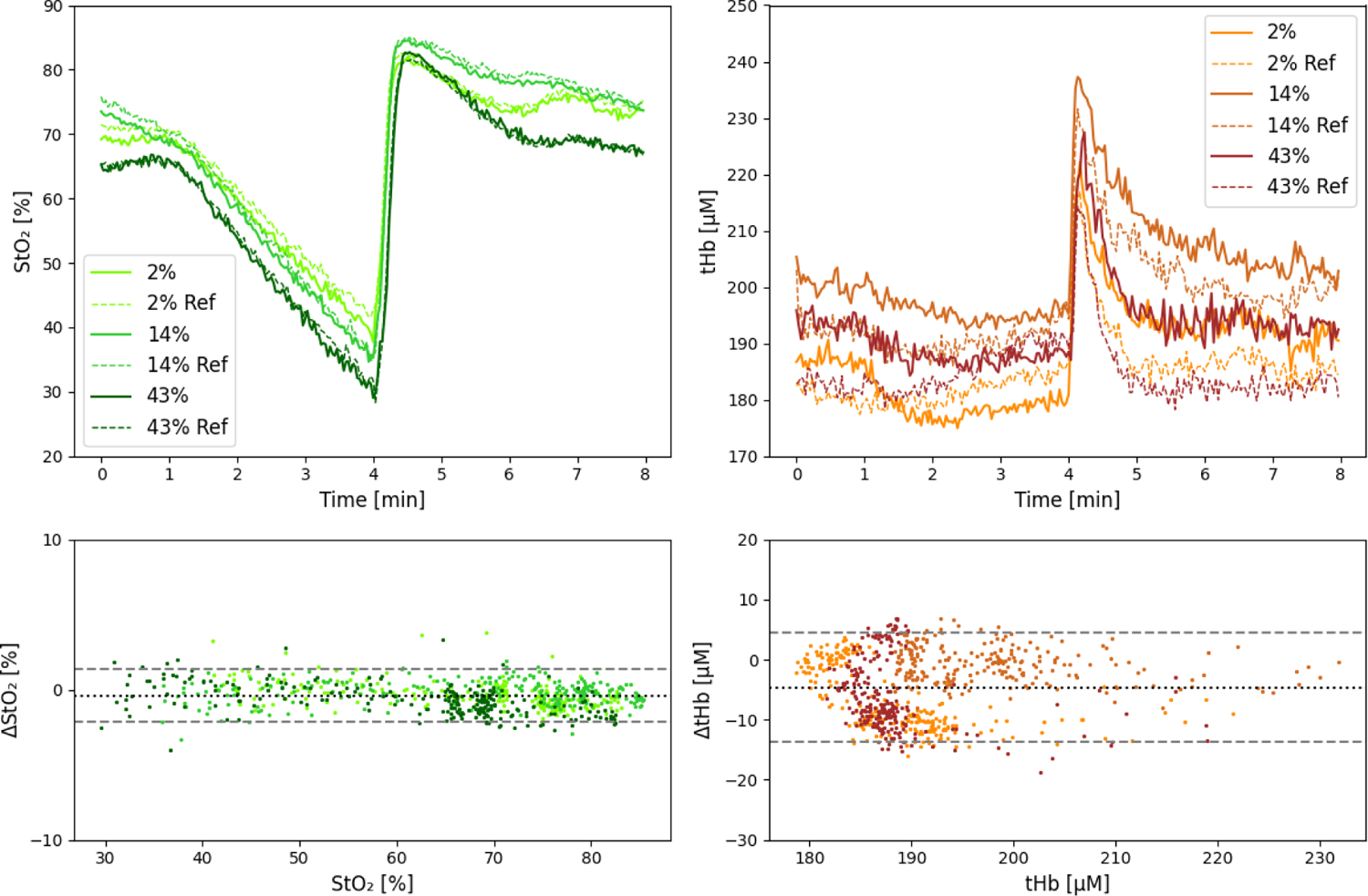
Time courses (top row) and Bland Altman plot (bottom row) for S_t_O_2_ (left column) and tHb (right column) during arterial occlusion with skin phantoms mimicking different skin pigmentation tone (M_f_ = 0%, 2%, 14%, 43%) for subject S08.

**Supplementary Figure 5:**
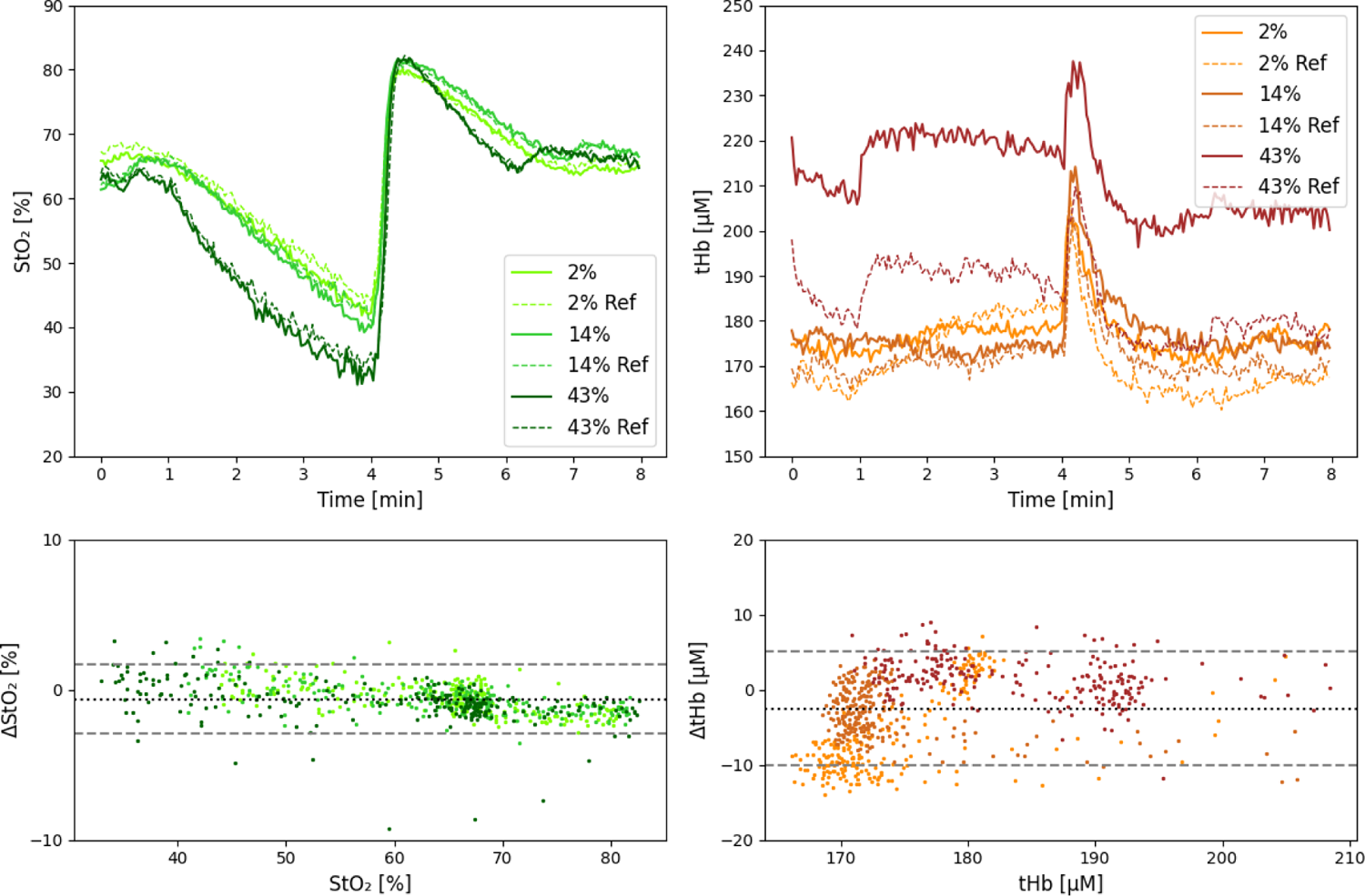
Time courses (top row) and Bland Altman plot (bottom row) for S_t_O_2_ (left column) and tHb (right column) during arterial occlusion with skin phantoms mimicking different skin pigmentation tone (M_f_ = 0%, 2%, 14%, 43%) for subject S09.

**Supplementary Figure 6.**
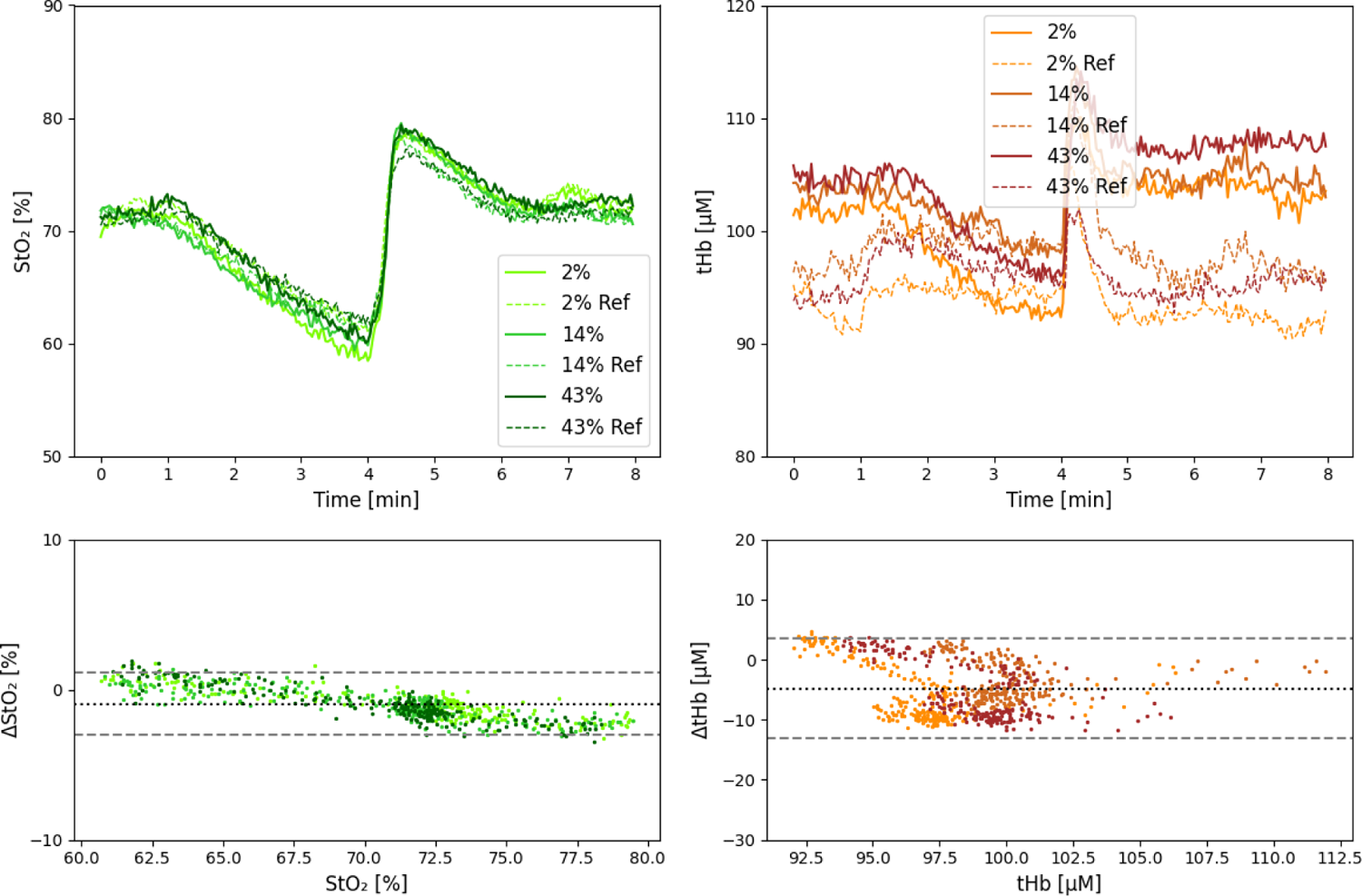
Time courses (top row) and Bland Altman plot (bottom row) for S_t_O_2_ (left column) and tHb (right column) during arterial occlusion with skin phantoms mimicking different skin pigmentation tone (M_f_ = 0%, 2%, 14%, 43%) for subject S10.

**Supplementary Figure 7:**
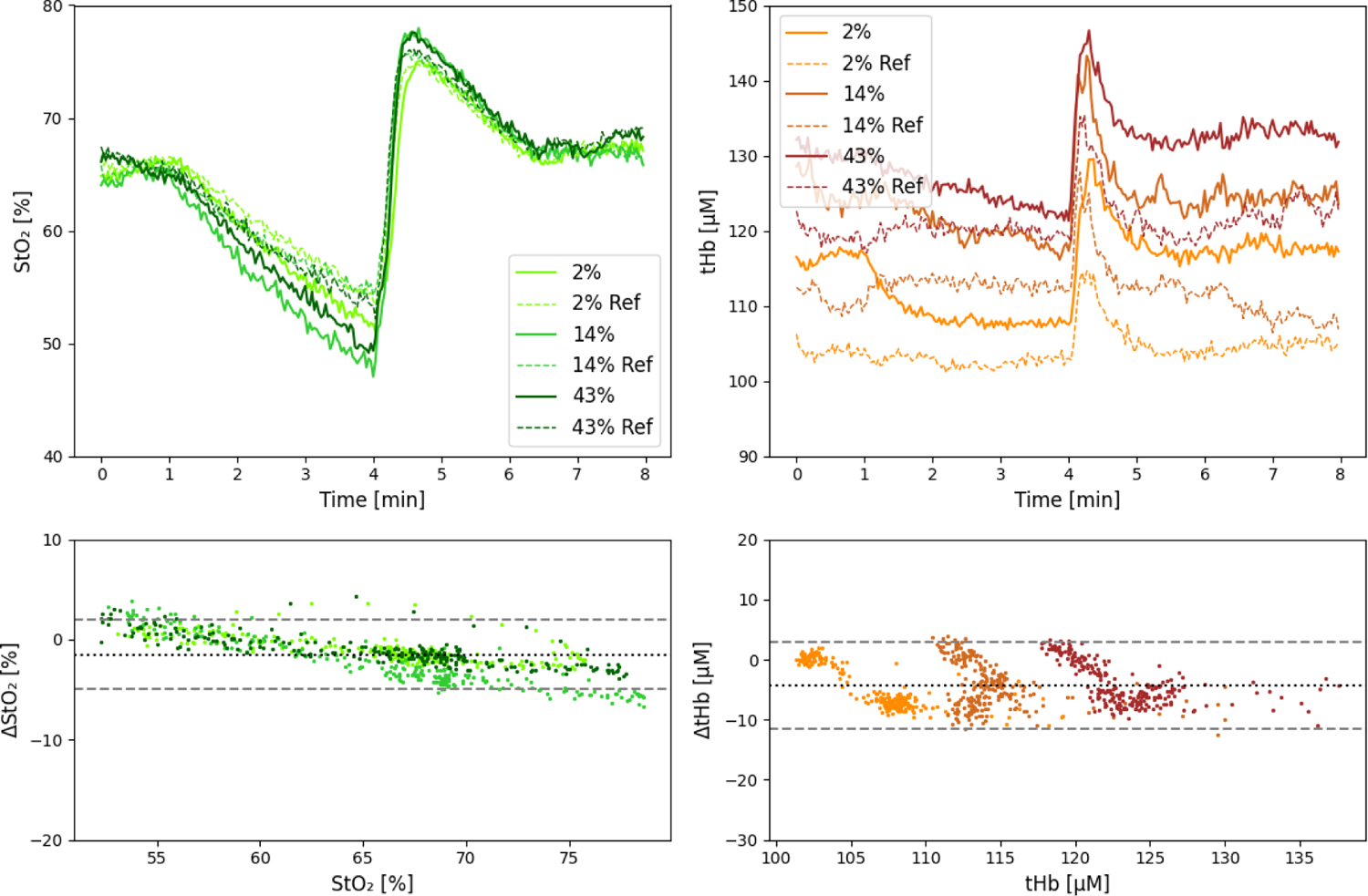
Time courses (top row) and Bland Altman plot (bottom row) for S_t_O_2_ (left column) and tHb (right column) during arterial occlusion with skin phantoms mimicking different skin pigmentation tone (M_f_ = 0%, 2%, 14%, 43%) for subject S12.

### S3 TD NIRS measurement setup

**Supplementary Figure 8:**
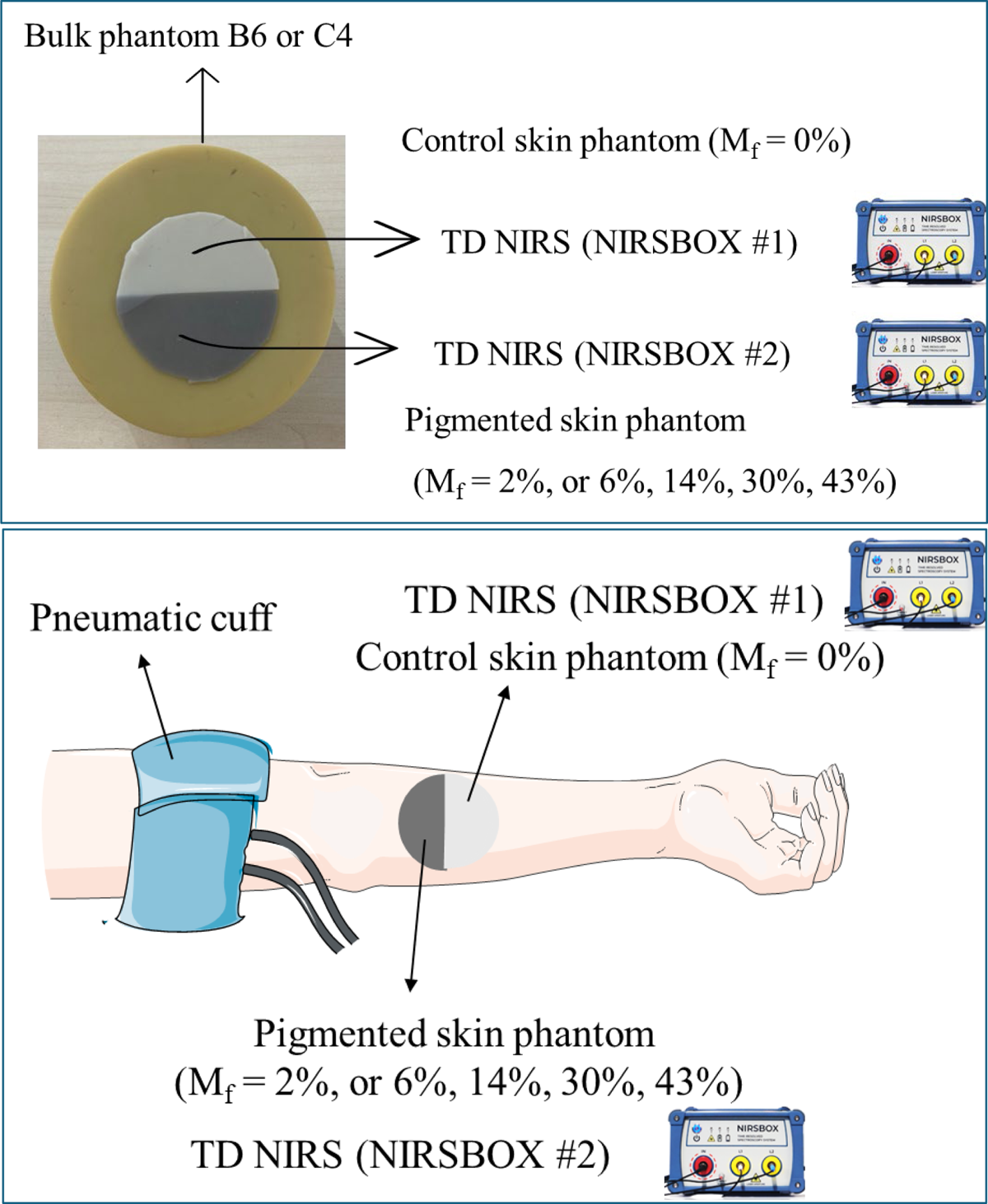
Setup used for TD NIRS measurements on bulk phantoms B6 and C4 (top) and for vascular occlusion test (bottom). Graphical abstract (arm and cuff occlusion) was drawn in part using images from Servier Medical Art (https://smart.servier.com/citation-sharing/) licensed under a Creative Commons Attribution CC-BY License.

